# A distinct circuit for biasing visual perceptual decisions and modulating superior colliculus activity through the mouse posterior striatum

**DOI:** 10.1101/2024.07.31.605853

**Authors:** Kara K. Cover, Kerry Elliott, Sarah M. Preuss, Richard J. Krauzlis

## Abstract

The basal ganglia play a key role in visual perceptual decisions. Despite being the primary target in the basal ganglia for inputs from the visual cortex, the posterior striatum’s (PS) involvement in visual perceptual behavior remains unknown in rodents. We reveal that the PS direct pathway is largely segregated from the dorsomedial striatum (DMS) direct pathway, the other major striatal target for visual cortex. We investigated the role of the PS in visual perceptual decisions by optogenetically stimulating striatal medium spiny neurons in the direct pathway (D1-MSNs) of mice performing a visual change-detection task. PS D1-MSN activation robustly biased visual decisions in a manner dependent on visual context, timing, and reward expectation. We examined the effects of PS and DMS direct pathway activation on neuronal activity in the superior colliculus (SC), a major output target of the basal ganglia. Activation of either direct pathway rapidly modulated SC neurons, but mostly targeted different SC neurons and had opposite effects. These results demonstrate that the PS in rodents provides an important route for controlling visual decisions, in parallel with the better known DMS, and with distinct anatomical and functional properties.

**Significance Statement:** The rodent posterior striatum (PS) is strongly innervated by visual cortex and thalamus, but its functional role in visual behavior has not been explored. We show that the PS initiates a direct pathway through the basal ganglia that is anatomically distinct from the more commonly studied dorsomedial striatum (DMS). Activating the PS direct pathway selectively biases decisions for expected, valued visual events. We also show that both DMS and PS direct pathways modulate neuronal activity in the superior colliculus, a structure critical for visual processing and sensorimotor function, and preferentially modulate collicular units with properties relevant in the visual detection task. These findings identify a distinct and novel circuit through the basal ganglia for controlling visually guided perceptual decisions.

## Introduction

The basal ganglia comprise a set of interconnected subcortical nuclei that govern a range of complex functions. The best-known functions are related to motor control. The striatum, the input nucleus of the basal ganglia, facilitates elements of action execution including movement (Freeze et al., 2013), goal-directed action selection (Tai et al., 2012) and orienting to valued sensory stimuli (Hikosaka et al., 2019). However, a growing body of evidence implicates the basal ganglia in aspects of perceptual decision-making that are distinct from motor control (Ding, 2023). Caudate neurons in primates exhibit spatially selective modulation in the absence of goal-directed actions (Arcizet and Krauzlis, 2018). Neurons in the caudate of primates (Ding and Gold, 2010) and dorsal striatum of rats (Yartsev et al., 2018) show activity consistent with sensory evidence accumulation. Moreover, deficits in sensory perception are noted in disorders of basal ganglia function, such as Parkinson’s (Permezel et al., 2023) and Huntington’s (Profant et al., 2017) diseases.

Sensory information is routed to the striatum through cortical and thalamic nuclei associated with somatosensory, visual, and auditory modalities (Hintiryan et al., 2016; Hunnicutt et al., 2016). The majority of studies on striatal involvement in visual perceptual behavior have focused on the dorsomedial striatum (DMS) in rodents and the head of the caudate nucleus in primates which receive input from visual cortex (Yeterian and Pandya, 1995; Khibnik et al., 2014) as well as visual thalamus (Smith and Parent, 1986; Kamishina et al., 2008). Dopamine D1-receptor expressing medium spiny neurons (D1-MSNs) in the striatum initiate the direct pathway, a disynaptic circuit that is believed to gate behaviors by disinhibiting basal ganglia targets in the thalamus and midbrain. Activating DMS D1-MSNs in mice biases visual decisions in a manner linked to the allocation of spatial attention (Wang et al., 2018; Wang and Krauzlis, 2020). Thus, several lines of evidence have established the direct pathway arising from the dorsal striatum as an important circuit for visual perceptual decisions.

The posterior striatum (PS) is also innervated by visual cortex and thalamus (Khibnik et al., 2014; Hunnicutt et al., 2016; Griggs et al., 2017; Lee et al., 2023). In the primate homologue, the tail of the caudate nucleus, neurons in the direct pathway encode visual cue value and bias the generation of saccades (Kim et al., 2017; Amita et al., 2019, 2020). In the rodent, a role for PS in visually guided behavior remains untested. PS D1-MSN signaling is causally involved in auditory decisions in mice (Guo et al., 2018; Chen et al., 2022). Also, dopamine axons in the PS respond to light (Menegas et al., 2018). It thus seems plausible that visual circuits through the PS could be a major contributor to visual perceptual function, but this has yet to be explored in rodents.

Here, we investigated a functional role for the PS direct pathway in visual perceptual decisions. We first characterized this circuit using anatomical methods and then tested its functional role by combining optogenetic PS or DMS D1-MSN manipulations with SC neuronal recordings in mice making visual decisions. We found that the direct pathway from the PS is largely segregated from that of the DMS and is more broadly and inclusively innervated by visual cortex. Activation of this PS direct pathway circuit biased performance in the visual change-detection task and modulated the activity of SC neurons. Whereas the predominate neuronal effect of PS or DMS D1-MSN activation was excitatory modulation of SC units, PS D1-MSN stimulation induced inhibitory effects more frequently than that of DMS. Together, these findings demonstrate that the PS provides a functionally significant and distinct route through the basal ganglia for visual perceptual decisions.

## Materials and Methods

### Subjects

Male and female C57BL/6J (wild-type; Jackson Laboratory, #000664) or Drd1a-Cre heterozygous (B6.FVB(Cg)-Tg(Drd1a-cre)EY217Gsat/Mmucd; MMRRC #034258-UCD) mice were housed a in temperature and humidity-controlled vivarium under a 12-hour reversed light/dark cycle (lights off at 0800 hours). All experimental procedures occurred during the dark cycle. Mice were housed with littermates (two to five per cage), except for those singly housed following head post and optic fiber implantation. Mice performing behavioral tasks were weighed and fed daily to maintain (at least) 85% *ad libitum* weight and the health status of each mouse was monitored daily throughout the study. These mice had free access to water, but their intake of dry food was controlled and were additionally given access to a nutritionally complete 8% soy-based infant formula (Similac, Abbott, IL). All experimental and animal husbandry procedures were approved by the NIH Institutional Animal Care and Use Committee (IACUC) and complied with the Public Health Service policy on the humane care and use of laboratory animals.

### Surgical procedures

Mice of at least 6 weeks of age were anesthetized with isoflurane (4% induction, 0.8-1.5% maintenance) and secured by a stereotaxic frame with ear bars. A feedback-controlled heating pad was used to maintain the body temperature at 37°C and ophthalmic ointment was applied to the eyes. Dexamethasone (1.6 mg/kg, s.c.) and meloxicam (2.0 mg/kg, s.c.) were administered to reduce inflammation and discomfort, respectively. Viruses were infused at a rate of 2 nl/s (50 nl/min maximum) using a microinjector (Nanoject III, Drummond Scientific, Broomall, PA).

For basal ganglia direct pathway tracing in Drd1-cre heterozygous mice, either AAV1-Ef1A-DIO-iChloc-2a-dsRed (NIDA Genetic Engineering and Viral Vector Core) or AAV1-Ef1A-DIO-eYFP (Addgene #27056-AAV1) was unilaterally injected in the DMS (200 nl; distance from bregma in mm, [anterior-posterior, AP] 0.0, [medial-lateral, ML] 2.0, [dorsal-ventral, DV] -2.1) and the other virus injected in the PS (100 nl at each depth; [AP] -1.5, [ML] 3.5, [DV] -2.5 and -3.1). The viruses injected in DMS and PS were counter-balanced across 3 mice.

For di-synaptic tracing of DMS and PS direct pathways through the SN in wild-type mice, either AAV1-hSyn-cre-WPRE-hGHpA (Addgene #105553-AAV1) or AAV1-Ef1a-flp (Addgene #55637-AAV1) was unilaterally injected in the DMS and the other virus in the PS (same volumes and coordinates as listed above), as well as Fluorogold (30 nl, 1.5%, Fluorochrome LLC, Denver, CO) to label the injection site. In the same mice, both AAV1-Ef1a-fDIO-mCherry (Addgene #114471-AAV1) and AAV1-Ef1a-DIO-eYFP (Addgene #27056-AAV1) were injected in the SNR (150nl each; [AP] -3.2, [ML] 1.6, [DV] -4.4). The viruses injected in DMS and PS were counter-balanced across 4 mice. For di-synaptic tracing of the DMS direct pathway through the EP (200 nl; [AP] - 1.25, [ML] 1.9, [DV] -4.3), a single set of viruses were used (either AAV1-cre and AAV1-DIO-eYFP, or AAV1-flp and AAV1-fDIO-mCherry), counter-balanced across 4 mice.

For retrograde tracing of projections to the striatum, Fluorogold (3%) was unilaterally injected in the DMS (80 nl; [AP] 0.0, [ML] 2.15, [DV] -2.1) of 2 mice or in the PS (40 nl at each depth; [AP] -1.5, [ML] 3.5, [DV] -2.5 and -2.95) of 2 mice.

For optogenetic manipulation of the direct pathway in mice performing behavioral tasks, AAV2-Ef1a-DIO-hChR2(H134R)-mCherry (University of North Carolina viral core) or a control virus AAV1-hSyn-DIO-mCherry (Addgene #50459-AAV1) was injected unilaterally in the DMS (250 nl; [AP] 0.0, [ML] 1.7, [DV] -2.1) and the PS (80 nl at each depth; [AP] -1.5, [ML] 3.25, [DV] -2.25 and -2.75). Additionally, optic fibers (200 µm core; Plexon, Dallas, TX) were inserted in the DMS ([AP] 0.0, [ML] 1.7, [DV] -1.7) and PS ([AP] -1.5, [ML] 3.25, [DV] -2.15) and a custom-designed titanium head post was secured to the skull using Metabond (Parkell Inc., Edgewood, NY). The skin wound edge was then closed with surgical adhesive and treated with antibiotic ointment. Mice assigned to behavioral testing began food restriction 18-20 days following surgery.

After mice were trained and tested on the visual detection task with optogenetic manipulation, a second surgery was performed to implant electrophysiology microwire bundles. Following the same anesthetic and analgesic procedure as detailed above, a small craniotomy was made to implant a drivable 16-wire microwire bundle (Innovative Neurophysiology, Durham, NC). The tip of the stainless-steel cannula of the bundle was implanted at [AP] -3.4 – -3.9, [ML] 0.8 – 1.2 and [DV] -0.2. The cranial opening was sealed with Vaseline petroleum jelly and the microwire bundle assembly was secured on the skull with Metabond.

Immunohistochemistry was performed to confirm virus expression and fiber placement. Animals were excluded from analysis for poor virus expression at target regions or viral expression in non-target regions (i.e. cortex).

### Immunohistochemistry

Mice were deeply anesthetized with isoflurane prior to transcardial perfusion with 0.1 M PBS, pH 7.3, followed by ice cold 4% paraformaldehyde in PBS. Brains were extracted and stored in 4% paraformaldehyde at 4°C. 50 µm coronal sections were cut using either a Campden Instruments 7000smz-2 vibrating microtome (Lafayette, IN) at room temperature or, following transfer to 20% sucrose solution, a Leica SM2010 R freezing microtome (Deer Park, IL). Rabbit anti-GFP (1:1000; G10362, Invitrogen, Carlsbad, CA) and donkey anti-rabbit conjugated to Alexa Fluor 488 (1:1000; A21206, Invitrogen) were used to amplify eYFP expression. Chicken anti-mCherry (1:2000; NBP2-25158, Novus Biologicals, Littleton, CO) and donkey anti-chicken conjugated to Alexa Fluor 594 (1:1000; 703-585-155, Jackson ImmunoResearch, West Grove, PA) were used to amplify dsRed and mCherry.

### Behavioral testing

#### Behavioral apparatus

Behavioral testing was conducted in a custom-built booth that displayed visual stimuli on the left- and right-hand sides of the head-fixed mouse that was coupled to their locomotion on a custom-built Styrofoam linear wheel(Krauzlis et al., 2020). A pair of LCD displays (VS17287; ViewSonic, Brea, CA) were positioned at a 45° angle from the midline of the animal so that the displays were centered on the right or left eye and subtended by ∼90° horizontal and ∼55° vertical of the visual hemifield. The viewing distance from the mouse to the screen was ∼27.5 cm. Sound-absorbent material lined the inside of the booth to reduce noise.

Experiments were controlled by a computer using a modified version the PLDAPS system (Eastman and Huk, 2012), consisting of a Datapixx peripheral (Vpixx Technologies, Saint-Bruno, QC, Canada) and the Psychophysics Toolbox (Brainard, 1997; Pelli, 1997) for MATLAB (MathWorks, Natick, MA), controlled by MATLAB-based routines run on a PC running Ubuntu 18.04 or on a Mac Pro (Apple, Cupertino, CA). The Datapixx device provided autonomously timed control of analog and digital inputs and outputs synchronized to the visual display stimuli. A spout delivering liquid rewards was placed near the mouth of the mouse and lick contacts on the spout were detected by a piezo sensor using custom electronics. ∼5 - 10 μL of 8% solution of infant formula were used as rewards for correct task performance and were delivered via a peristaltic pump.

#### Visual detection tasks

The detection task was modified from our previously published study (Wang et al., 2018). Animals were run in daily sessions producing 153 to 421 trials, with data pooled across sessions. Experiments were organized in blocks of randomly shuffled trials. Each trial consisted of a sequence of epochs that were defined by the stimuli presented on the visual displays. Epoch durations were determined by either the randomized timing of the epoch or the time that it took for the mouse to travel a randomized distance on the wheel. Mice were trained to detect a change in the orientation of a Gabor patch visual stimulus, which the mouse reported by contacting the lick spout to receive a reward.

The trial sequence of the primary detection task consisted of four epochs (Figure 5A). The average luminance across each visual display in all epochs was 4 – 8 cd/m^2^. In the first epoch (“Pink noise”), the uniform gray of the intertrial interval was changed to pink visual noise with a root mean square contrast of 3.3%. In the second epoch (“Gabor patch cue”) a vertically oriented Gabor patch was added to the pink noise, centered on either the left or right visual display. The sinusoidal grating of the Gabor patch (93.7% Michelson contrast) had a spatial frequency of 0.1 cycles per degree and was modulated by a Gaussian envelope with a full width at half maximum of 18° (standard deviation = 7.5°). The phase of the grating was incremented in proportion to the wheel rotation and updated on every monitor refresh so that the sinusoidal pattern displaced on the screen matched the distance that the mouse traveled on the wheel. The Gabor patch on the left (right) drifted leftward (rightward), consistent with optic flow during locomotion. The third epoch (“Delay epoch”) consisted of vertical Gabor patches appearing in both left and right visual displays. The stimuli in the fourth epoch depended on whether the trial was a “Change” or “Catch” condition; the two trial types were equally likely and randomly interleaved within a block. On Change trials, the Gabor patch on the same side as the patch present in second epoch changed its orientation (10-13°, depending on the mouse). The direction of the orientation change was clockwise for the left patch and counterclockwise for the right patch. On catch no-change trials, neither Gabor patch changed orientation, such that the epoch proceeded as a seamless extension of the previous delay epoch.

Mice were to lick the spout upon detection of a change in the orientation of the Gabor patch and otherwise withhold licking. Mice were required to lick within a 550 ms response window starting 220 ms after the onset of the orientation change to score a “hit” and receive a reward. If the mouse failed to lick within the window, the trial was scored as a “miss” and no reward was given. On catch trials, if the mouse licked with the same response window aligned on the no-change epoch onset, the trial was scored as a “false alarm” and was used to calculate lick probability rates across trial types. If the mouse correctly withheld licks during this window, the trial extended to include an additional “Safety” epoch in which the initially appearing Gabor underwent a 30° suprathreshold orientation change and the mouse could receive a reward by licking within a comparable response window. This safety epoch was intended to maintain motivation by rewarding mice for correct behavior without violating the task rule that they should lick only for orientation changes. False alarms or two or more premature licks before the response window resulted in termination of the trial and a three second timeout.

In half of the trials, 465 nm blue light was delivered unilaterally to either the PS or DMS using an LED system (Plexon). For half of these trials, light was delivered during the delay epoch (“Early delay”) for 150 ms, starting 200 ms after the onset of the epoch. For the remaining 50% of stimulation trials, light was delivered during the fourth epoch, either during catch (“Late delay”) or change trials with equal frequency, for 150 ms, starting 50 ms after the onset of the epoch. Light power was set at a fixed intensity per striatal target per mouse to induce a non-saturating increase in lick probability (i.e., less that 100%) and ranged from 0.05 to 1.15 mW across the ChR2-expressing cohort. Similar light intensities were used for mice expressing the control mCherry fluorophore.

Trials were organized into alternating 40-trial blocks in which the orientation change occurred for either the left or right Gabor patch. Each block had an equal number of trials with and without orientation changes, and with and without optogenetic light delivery, all randomly shuffled. Each session started with a mini-block of 20 trials without light delivery trials which were excluded from behavioral analysis.

A “sequence arrested” control experiment consisted of blocks of 54 trials, 40 of which followed the standard four-epoch trial sequence: 20 with an orientation change (half of which were paired with optogenetic stimulation during the change) and 20 no-change catch trials (half of which were paired with optogenetic stimulation during the late delay catch epoch). Two additional trial types were introduced in which the visual sequence of epochs stopped once the trial reached a particular epoch: seven stopped at the single Gabor patch and seven stopped at the pink noise. These sequence-arrested trials had the same durations and matched timing for optogenetic light delivery as the standard four-epoch trial sequence. We recorded 2-3 sessions per mouse with PS D1-MSN stimulation and pooled across sessions.

To test the interaction of PS D1-MSN stimulation on unreinforced change detection responses, we conducted a version of the detection task with four trial types: catch and change trials (50% of each), with stimulation during the change or late delay on half of each. Instead of blocks of trials switching between left- and right-side orientation changes, we only presented trials with the Gabor patch cue and orientation change occurring in the visual field contralateral to the hemisphere receiving optogenetic stimulation. After a 60-trial block in which correct change detection was rewarded with soymilk rewards, the remaining trials were not reinforced, and the session was terminated after the mouse extinguished lick responses. We recorded 2-3 sessions per mouse and pooled across sessions and mice for subsequent analysis.

### In vivo electrophysiology

In the cohort of Drd1a-cre mice expressing DIO-ChR2-mCherry and optic fibers in PS and DMS, we obtained electrophysiological recordings from isolated neurons in the SC while mice performed the orientation change detection task with optogenetic activation of PS and DMS D1-MSNs in the same hemisphere. The extracellular activity of SC neurons was recorded with microwire bundles. Electrophysiological signals were acquired through an RZ5D processor (Tucker-Davis Technologies, Alachua, FL) with spikes filtered between 0.3 and 5 kHz and sampled at 25 kHz. Single units were sorted offline using KiloSort (Pachitariu et al., 2016). The SC surface was determined based on the first visual responses detected while advancing the microwire bundle. Sessions alternated between optogenetic stimulation of either PS or DMS D1-MSNs.

Visual receptive field maps were obtained at the end of every recording session, during which white circular disks (118.5 cd/m^2^) of 10° in diameter were flashed against a gray background (7.2 cd/m^2^) in the visual display contralateral to the recording side. We sampled visual locations pseudo-randomly drawn from a 3 x 7 isotropic grid that extended from -25° to 25° in elevation and 0° to 90° in azimuth of the contralateral visual field. An individual trial consisted of 8 consecutive 250 ms flashes, which flash followed by a 250 ms blank interval. At least 9 flash repetitions for each grid location were presented in each mapping session. Receptive fields of individual neurons were estimated from the mean spike counts 50 – 150 ms after the flash onset in each grid location, after subtracting baseline activity. The baseline in each trial was defined as the mean spike count within the 100 ms period before the presentation of the first flash. Baseline-subtracted mean spike counts were linearly interpolated between grid locations with 1° resolution using the *scatteredInterpolant* function in Matlab.

Following the last recording session, we performed electrolytic lesions to aid in histological reconstruction of bundle location within the SC. Under isoflurane anesthesia, 80 µA current was delivered for 7-10 s to two wires per bundle. Mice were administered meloxicam and perfused one week later.

### Eye movement monitoring

To verify that changes in SC neuronal activity were not simply due to eye movements, a 120Hz video eye tracking system (ETL-200, ISCAN, Woburn, MA) was used to monitor eye position and pupil size of head-fixed mice during electrophysiology experiments with optogenetic manipulation. The frequency of eye movements occurring in trials with and without optogenetic D1-MSN stimulation per stimulation site, per mouse, was statistically tested with chi-square tests.

### Quantification

#### Behavioral analyses

Lick probability was quantified separately for contralaterally and ipsilaterally cued trials and light delivery and no light delivery trials. Probability was calculated from the number of trials in which there were any licks within a 750 ms window starting at onset of light delivery for light delivery trials and a matched time window for non-light delivery trials (i.e. 200 ms to 950 ms from onset of the early delay epoch and 50 ms to 800 ms from the onset of the late delay catch epoch) divided by the total number of trials within the assessed trial condition, per mouse. Reaction time was calculated from the first lick following the orientation change per trial condition per mouse.

To compare behavioral extinction rates we separately fit binary responses (1 = hit, 0 = miss) to the orientation change for non-stimulation and PS D1-MSN stimulation trials using a four-parameter logistic regression model (Subramanian et al., 2023):

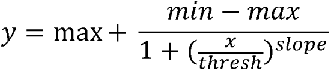

In which *x* is the trial number relative to reward non-reinforcement (1 = the first unrewarded trial), *y* is the value of the function, *min* is the minimal value of the curve, *max* is the value of y at the lowest value of x, *thresh* is the inflection point, and *slope* is the slope of the function. The four parameter terms were fit to binary data by minimizing mean squared error. For each data set, 95% CIs for the four parameters were estimated using a standard bootstrap procedure with 1000 random draws. Significant differences between the fitted curves for non-stimulation and PS stimulation trials were assessed with z-tests for specific values of y (probability of a hit).

#### Neuronal analyses

Firing rates for each neuron were summarized as peri-stimulus time histograms (PSTHs) by counting spikes within 20 ms non-overlapping bins aligned to key task events. When pooling firing rates across neurons, these spike counts were converted to normalized z-score values by subtracting the mean spike count from the PSTH and dividing by the standard deviation, using the mean and standard deviation of the spike counts calculated from the 20 ms bins within the trials across the entire recording session. Population PSTHs presented in figures were generated from spike counts in 20 ms bins sliding in 1 ms intervals and normalized to z-scores, with error bars representing standard error.

Phasic responses to the Gabor patch cue were tested by either Wilcoxon signed rank test on spike counts in the interval 50 – 150 ms after Gabor patch cue epoch onset to spike counts -100 – 0 ms before Gabor patch cue epoch onset on contralaterally cued trials (relative to the recorded hemisphere) or Wilcoxon rank sum test on spike counts in the interval 50 – 150 ms after Gabor patch cue epoch onset on contralaterally cued trials and spike counts in the same interval for ipsilaterally cued trials. Delay-related activity was tested by Wilcoxon signed rank test on spike counts in the interval 200 – 300 ms after delay epoch onset to spike counts in the final 100 ms of the same epoch. Spatial cue -related modulation was tested by unpaired t-test of spike counts in the last 200 ms of the delay epoch on contralaterally cued trials to that of ipsilaterally cued trials. Phasic response to the orientation change was tested by Wilcoxon signed rank test of spike counts in the interval 50 – 150 ms after change epoch onset to -100 – 0 ms before change epoch onset on contralaterally cued trials. Lick related activity was tested by Wilcoxon signed rank test of spike counts in the interval -50 – 50 ms relative to first detection lick during the change epoch to spike counts in the interval -100 to 0 ms prior to change epoch onset. Significant differences in the proportions of units exhibiting task-related properties were assessed by chi-square tests.

SC neuronal responses to optogenetic direct pathway stimulation were tested by Wilcoxon signed rank test on spike counts in the interval 0 – 150 ms relative to light delivery to spike counts in a time-matched interval on non-stimulation trials and manually validated. Responses in each stimulation epoch (early and late delay, contralateral and ipsilaterally cued trials) were separately assessed for optogenetic modulation.

#### Histological analysis

Fluorescence density maps were generated as followed. Micrographs were converted to 8-bit and underwent contrast stretching to expand fluorescence intensity range from 0 to 255 using ImageJ Fiji (v. 1.54k). Images were loaded into Python (v. 3.11) and pixel intensity was converted to a z-score value calculated from all pixel values within the entire image of the micrographs of the SN or within the boundaries of the SC. A 50 um grid was applied to each image. A threshold of 1 standard deviation was then applied and for each 50 by 50 µm square, the pixels above this threshold were summed and divided by the total number of pixels, then multiplied by 100 to produce a percentage of pixels above threshold. This percentage is represented in grayscale, ranging from 0 (white) to 100% (black).

### Experimental design and statistical analyses

Sample sizes were determined based on publication standards. Studies were completed across multiple cohorts that produced homogenous experimental findings. For behavioral experiments, animals were randomly assigned to control and experimental groups. Group assignment was also balanced to ensure equal numbers of males and females.

Statistical tests were performed with Matlab (R2022a and R2023a) or GraphPad Prism 10 (Boston, MA) software. Error bars in figures represent 95% CI or standard error of the mean (s.e.m.) as indicated in figure legends. 95% CIs in figure plots were generated with the *binofit* Matlab function. The value of N represents the number of animals and n represents the number of units. All statistical tests were conducted two-sided, when applicable. Post-hoc Holm-Ŝidàk tests were performed to correct for multiple comparisons for ANOVAs; significant test results are indicated in the figures. All asterisks in figures indicate: * p<0.05, ** p<0.01, *** p<0.001, and **** p<0.0001.

Effect sizes were calculated for significant statistical comparisons. The eta squared (η2=SS_Effect_/SS_Total_) was calculated for 1-way ANOVA and the partial eta squared (partial η2=SS_Effect_/(SS_Error_+ SS_Effect_)) was calculated for N-way ANOVA main effects and interactions, where SS = sum of squares. Cohen’s d was calculated for post-hoc pairwise comparisons (d = mean difference/pooled standard deviation). For comparisons assessed with Chi-square tests, Cohen’s d was calculated from the chi square test statistic and sample sizes (n), as follows: 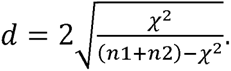

## Results

### The posterior striatum provides a unique basal ganglia direct pathway for visual signals

The rodent striatum is canonically divided into dorsomedial, dorsolateral, ventral, and posterior (or tail) subregions. D1-MSNs populate all regions of the striatum and provide the origin of the direct pathway of the basal ganglia. We sought to examine whether the direct pathway originating from the PS is anatomically distinct or shared with that of the DMS. To do this, we expressed cre-dependent anterograde fluorophore - expressing viruses in the PS and DMS of Drd1a-cre heterozygous mice (Figure 1). We found that DMS labeled D1-MSNs terminated in the ventral medial segment of the substantia nigra (SN) and in the entopeduncular nucleus (EP; homolog to the primate internal segment of the globus pallidus). In contrast, PS labeled D1-MSNs innervated the dorsal lateral region of the SN, with no labeling in the EP.

**Figure 1.**
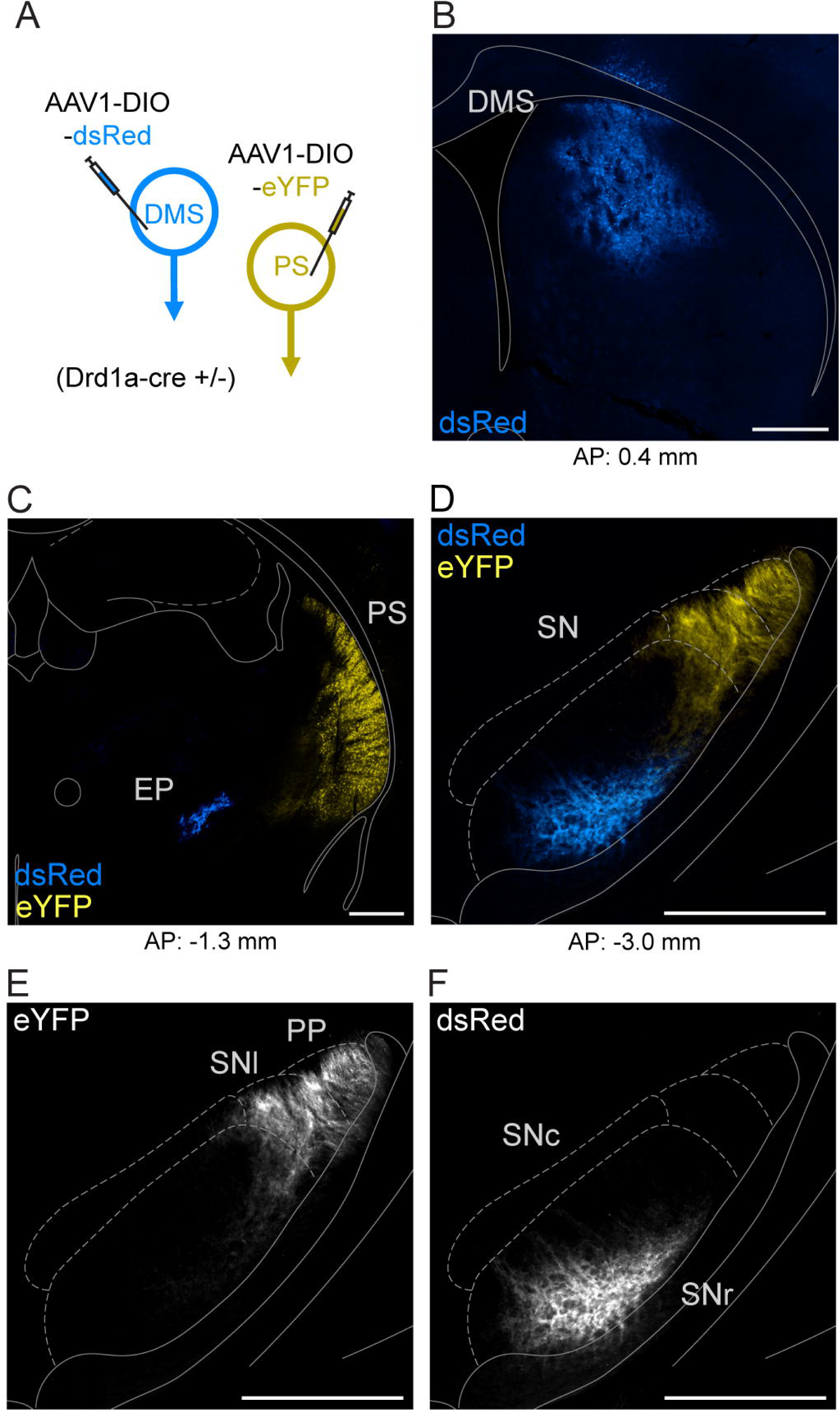
PS and DMS D1-MSNs innervate different basal ganglia output nuclei. A. Viral strategy to dual-label posterior (PS) and dorsomedial (DMS) D1-MSNs using cre-dependent anterograde viruses in D1-cre transgenic mice. B. Injection site of eYFP-expressing virus in DMS (blue). C. Injection site of dsRed-expressing virus in PS (yellow); axonal labeling of DMS-originating D1-MSNs in the entopeduncular nucleus (EP; blue). D-F. Axonal labeling of PS and DMS -originating D1-MSNs in the substantia nigra (SN). Anatomical boundaries approximated from Franklin & Paxinos Atlas (2008). Scale bars: 500 µm. Abbreviations: AP (anterior-posterior), PP (peripeduncular nucleus), SNl (substantia nigra pars lateralis), SNr (substantia nigra pars reticulata).

We then used a viral strategy to selectively label the neurons in the SN that are postsynaptic to PS and DMS D1-MSNs. We injected anterograde trans-synaptic AAV1 viruses in the PS and DMS of wild-type mice that delivered cre or flp recombinases to all efferent cells. We then co-injected cre- and flp-dependent non-trans-synaptic fluorophore-expressing viruses in the SN to locally label all recombinase-expressing cells putatively postsynaptic to D1-MSNs (Figure 2). This strategy selectively targets D1-MSNs because D2-MSNs do not project to SN (Lein et al., 2007; Kreitzer and Malenka, 2008; Allen Institute for Brain Science, 2025a, 2025b). We found neurons targeted by PS D1-MSNs were largely confined to the substantia nigra pars lateralis (SNl) and adjacent peripeduncular nucleus (PP) (Figure 2B). In contrast to the PS, neurons targeted by DMS D1-MSNs resided nearly exclusively in the substantia nigra pars reticulata (SNr) and in the overlying substantia nigra pars compacta (SNc) (Figure 2C). The presence of labeled cells in the SNc, a region that appeared to be sparsely innervated along the SNr border from DMS D1-MSN axons (Figure 1F), may be a result of retrograde labeling of striatal afferents which is noted to occur in recombinase-expressing AAV1 viruses (Zingg et al., 2020). Density mapping of fluorescent expression further demonstrated a general segregation of neuronal populations innervated by either PS or DMS (Figure 2D-E). By following the axonal expression of the neurons targeted by the PS and DMS D1-MSNs, we found a mixture of both exclusive and shared output targets in the diencephalon and midbrain (Figure 2F-L, O-P). To characterize the second direct pathway originating from the DMS and determine its efferent targets, in a separate set of cases we applied this di-synaptic labeling approach to confirm the projection from DMS D1-MSNs to neurons in the EP, with axons that innervate several midbrain nuclei and the habenula (Figure 3).

**Figure 2.**
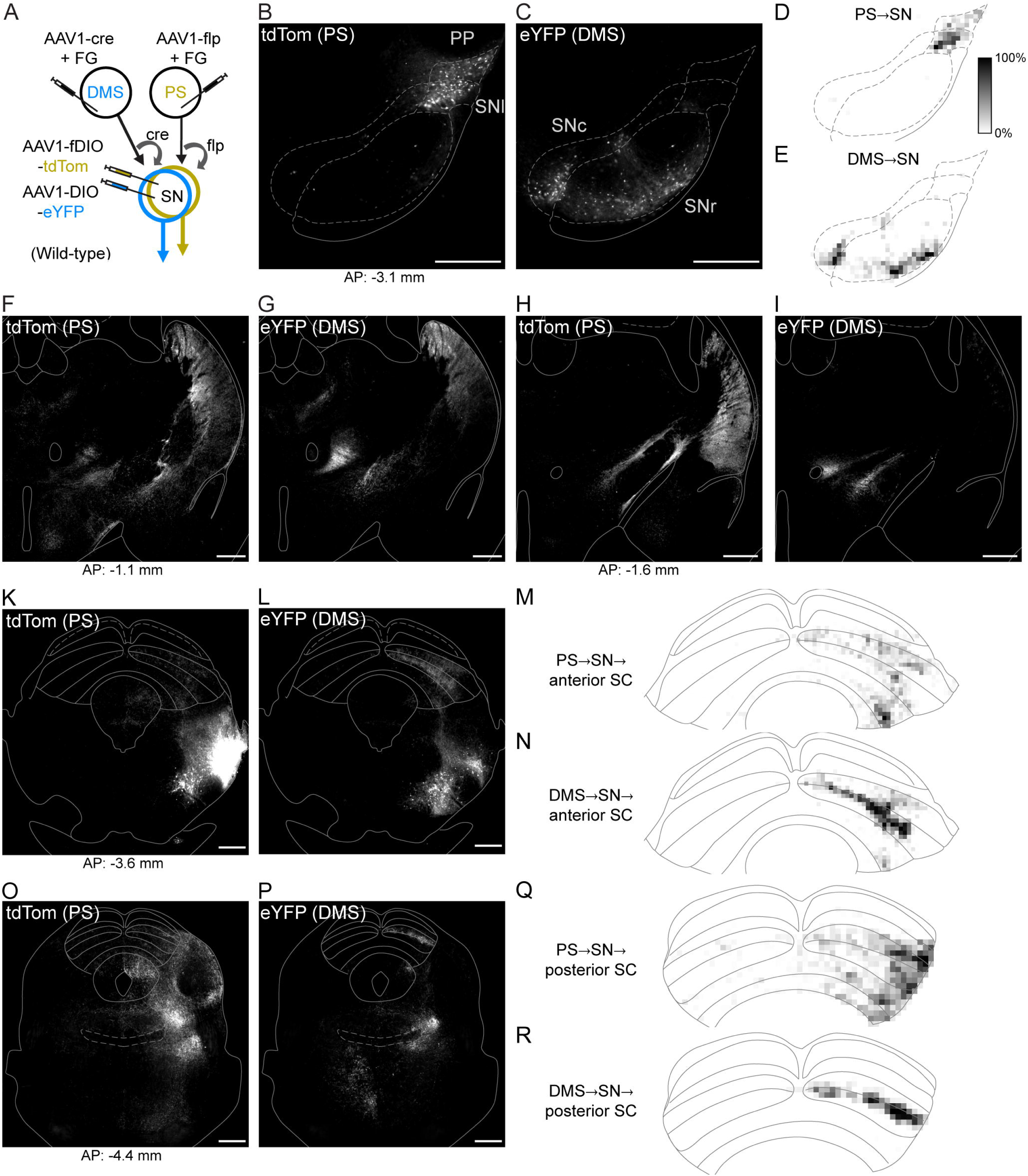
PS and DMS direct pathway circuits through the nigra are anatomically distinct but have shared output targets. A. Viral strategy to dual label PS and DMS -originating SN disynaptic pathways. B. Expression of DMS-innervated eYFP-labeled cells in the SNc and SNr. C. Expression of PS-innervated tdTom-labeled cells in the SNl and PP. D-E. Fluorescence density maps for cells and associated axons postsynaptic to the PS (D) and DMS (E). Grayscale represents the % of pixels within a 50 by 50 um area that exceed one standard deviation of z-scored fluorescent intensity (black = 100%). F-L, O-P. Terminal expression of PS→SN (tdTom) and DMS→SN (eYFP) cells in thalamus and midbrain. M-N, Q-R. Density maps of PS or DMS -initiating direct pathway terminal innervation of the superior colliculus (SC). Scale bars: 500 µm. Abbreviations: FG (Fluorogold).

**Figure 3.**
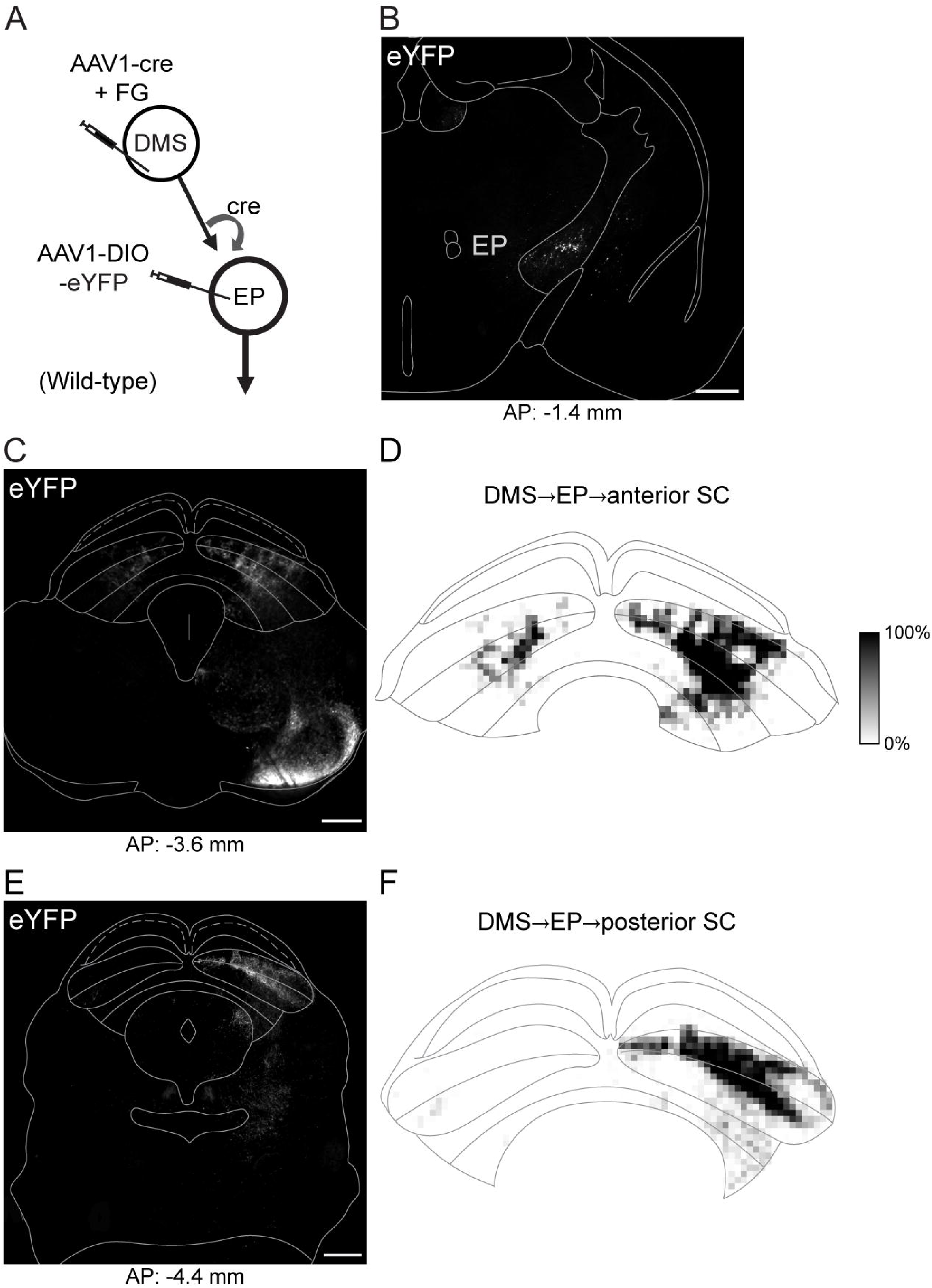
The DMS direct pathway through the EP innervates the SC. A. Viral strategy to label the DMS -originating EP disynaptic pathway. B. Labeled EP cells innervated by the DMS. C,E. Terminal expression of DMS→EP cells in the midbrain. D,F. Fluorescence density maps of terminal expression in the SC. Scale bars: 500 µm.

Although direct pathways originating from both DMS and PS innervate the SC, we observed striking differences. DMS-originating terminals showed concentrated innervation of SC intermediate layers (Figure 2N,R; Figure 3D-F), whereas PS-originating terminals exhibited diffuse innervation across all layers with relatively greater expression laterally and in more caudal sections of the SC (Figure 2M,Q).

Having established that D1-MSNs in the PS and DMS give rise to anatomically distinct pathways through the basal ganglia, we next compared the extent of visual cortical innervation to these two striatal subregions. Prior work indicates that the PS and DMS are the exclusive striatal targets of visual cortex (Hunnicutt et al., 2016). To identify the regions of visual cortex that project to these striatal regions, we injected the retrograde tracer Fluorogold unilaterally in either the PS or DMS of wild-type mice. We observed strong labeling of PS-projecting cells in what we identified as layers 5 and 6 across the entire rostral-caudal axis of the visual cortex, both in primary and secondary visual areas (Figure 4A-D). In contrast, we found strong labeling of DMS-projecting cells in layer 5 in rostral visual cortex but the labeling was weaker in more caudal sections, especially for primary visual cortex (Figure 4E-H). We also observed weak labeling of putative layer 3 cells for both striatal regions (PS, Figure 4B; DMS, Figure 4F). These results agree with similar findings in rats demonstrating stronger innervation from the visual cortex and thalamus to the PS than to the DMS (Jiang and Kim, 2018).

**Figure 4.**
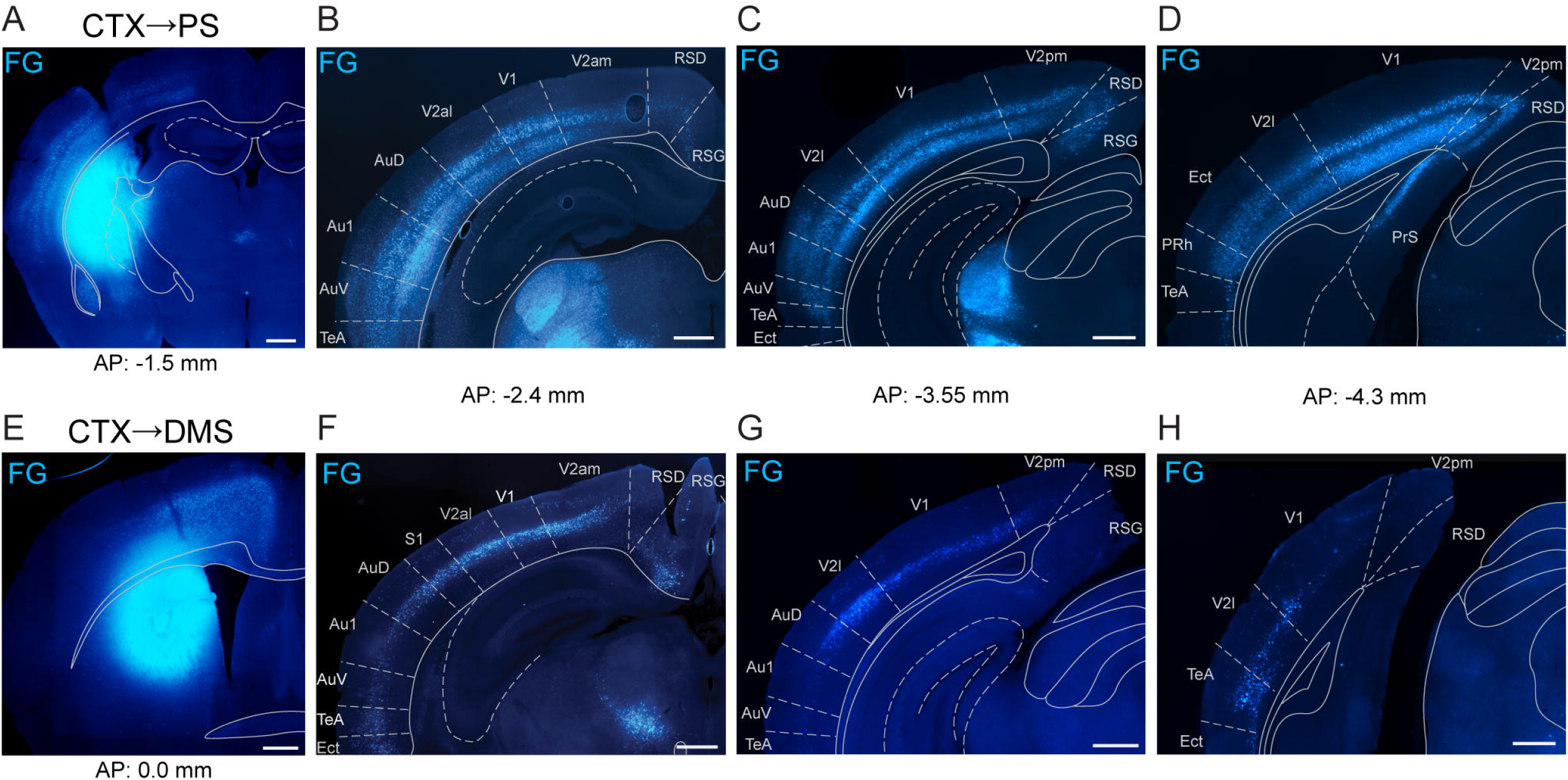
Visual cortex innervates PS more broadly and inclusively than DMS. A. Injection site of Fluorogold (FG) retrograde tracer in the PS. B-D. PS projecting FG-labeled neurons in visual cortex. E. Injection site of FG in the DMS. F-H. DMS projecting FG-labeled neurons in visual cortex. Cortical divisions approximated from Franklin & Paxinos and Allen Brain Atlases. Scale bars: 500 µm. Abbreviations: Au1 (primary auditory cortex), AuD (dorsal secondary auditory cortex), AuV (ventral secondary auditory cortex), CTX (cortex), Ect (ectorhinal cortex), PrS (presubiculum), RSD (retrosplenial dysgranular cortex), RSG (retrosplenial granular cortex), TeA (temporal association cortex), V1 (primary visual cortex), V2al (anterolateral secondary visual cortex), V2am (anteromedial secondary visual cortex), V2l (lateral secondary visual cortex), V2pm (posteromedial secondary visual cortex).

Collectively these experiments establish the PS as a distinct region within the striatum for influencing visual behavior. Unlike the DMS, the PS is strongly innervated by multiple layers through the entirety of visual cortex. By routing through the SNl and PP, the PS initiates a direct pathway largely segregated from that of the DMS. Lastly, we identified three distinct circuits by which the DMS and PS may modulate downstream targets, including the superior colliculus.

### Stimulation of the posterior striatum biases visual decisions

Our neuroanatomical results establish that the PS direct pathway provides a major route for visual signals through the basal ganglia that is distinct from the circuits emanating from the DMS. We next probed for functional roles of this striatal region by optogenetically activating PS D1-MSNs in mice performing a spatially cued orientation change-detection task (Wang and Krauzlis, 2018; Wang et al., 2018). Briefly, head-fixed mice were presented a vertical Gabor patch cue in one visual field that drifted at a rate coupled to running speed (Figure 5A). As the trial proceeded, a second Gabor patch appeared in the opposite visual field. After a variable time, the patch that had appeared first either changed orientation or remained vertical (‘catch’; 50% occurrence for each). Mice were trained to report their detection of the orientation change by licking a centrally placed spout to receive a soymilk reward and otherwise suppressed licks during other trial epochs (Figure 5B). Sessions were run in 40-trial blocks in which the orientation changed occurred exclusively in one visual field (left or right) before switching to the other location.

**Figure 5.**
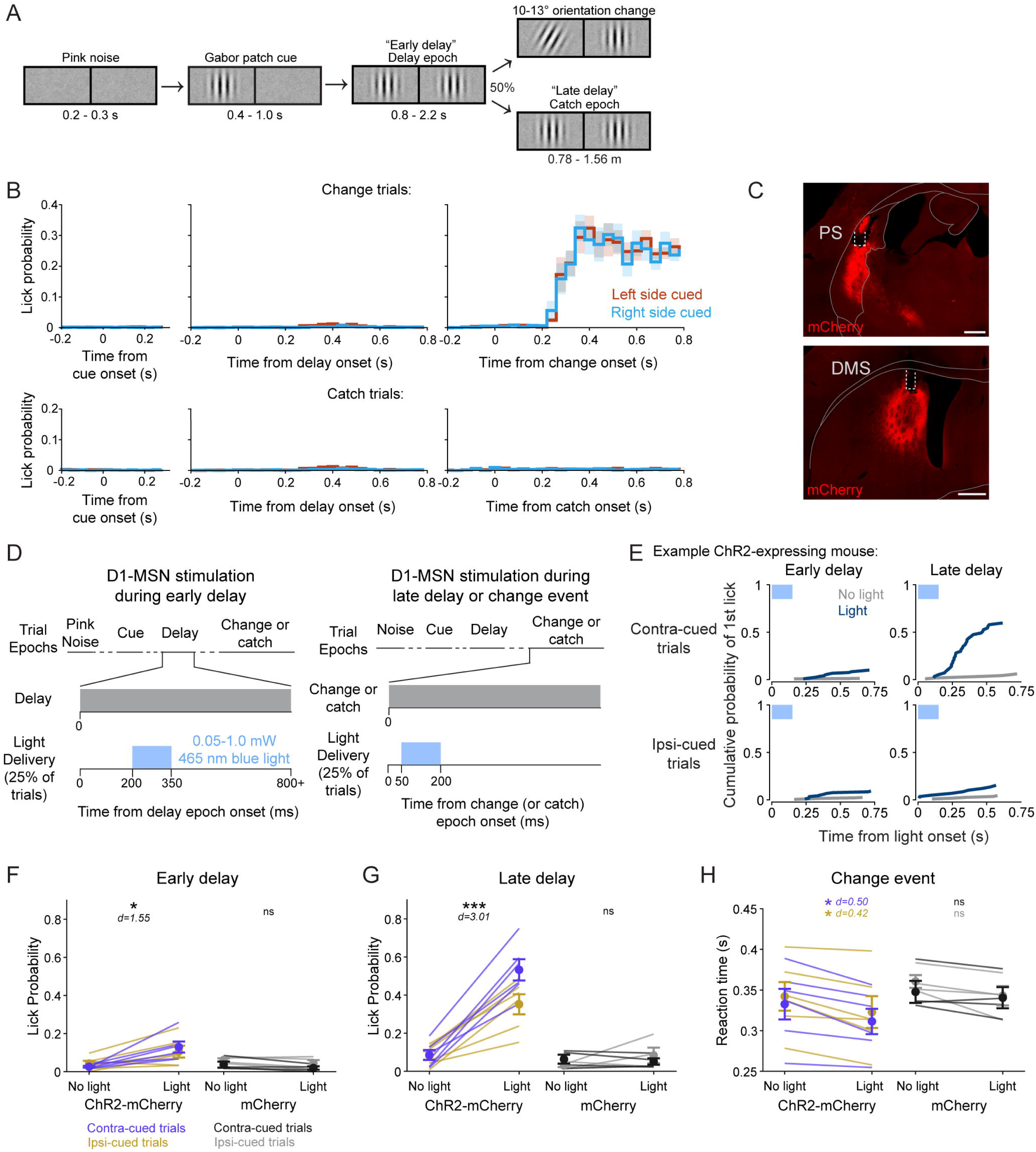
PS direct pathway activation biases visual decisions. A. Schematic of spatially cued orientation change-detection task trial structure and left and right visual displays. B. Peri-stimulus event histograms (PSTHs) of mean lick rates in 40-ms bins aligned to trial epoch onsets for change (top) and catch (bottom) trials (N=10). C. Representative expression of cre-dependent Channelrhodopsin (ChR2-mCherry) and optic fiber placement in the PS (top) and DMS (bottom) of a Drd1a-cre mouse. D. Schematic of light delivery to PS or DMS during the early delay (left) and late delay or change epochs (right). E. Cumulative lick probabilities from an example ChR2-expressing Drd1a-cre mouse. The first lick following PS light delivery onset (blue) or at a matched time on non-stimulation trials (gray) was quantified for the early delay epoch (left column) and late delay epoch (right column), separately for contralaterally cued trials (top row) and ipsilaterally cued trials (bottom row). F. Light delivery to the PS during the early delay epoch produced a small but significant increase in lick probability for ChR2-mCherry (purple and gold; N = 6) but not mCherry control (black and gray; N = 4) mice. G. PS light delivery during the late delay epoch significantly increased lick probability for ChR2-mCherry but not mCherry control mice. H. Light delivery during the change event reduced the median response latency for ChR2-mCherry but not mCherry control mice. Scale bars: 500 µm. Four-way repeated measures ANOVA (F, G); three-way repeated measures ANOVA (H). Error bars represent standard error of the mean (s.e.m.). d = Cohen’s d effect size. Abbreviations: ns (not significant).

We unilaterally stimulated channelrhodopsin (ChR2)-mCherry -expressing PS or DMS D1-MSNs in Drd1a-cre mice during one of three epochs of the task (Figure 5C-D). On 50% of the trials, 150 ms blue light was delivered either shortly after the appearance of the second Gabor patch (‘early delay’), during the change event (‘change’), or at a time-matched period on catch trials (‘late delay’). The visual conditions were identical between the early delay and late delay conditions (the presence of two vertical Gabor patches). We assessed lick responses within 750 ms of light delivery onset (or an equivalent time window on no-stimulation trials).

Light delivery to the PS during the early delay produced a small but significant increase in lick probability (‘yes’ response) in ChR2-mCherry-expressing mice (N = 6) compared to non-stimulation trials (Figure 5E-F; Table 1). Delivering light to the PS of mice expressing a control mCherry fluorophore (mCherry; N = 4) did not result in significant changes in lick probability (Figure 5F; Table 1). Delivering light during the late delay (in the absence of an orientation change) induced a larger significant increase in lick probability for ChR2-mCherry mice (Figure 5E,G, Table 1). Again, the control mCherry mice showed no effect (Figure 5G; Table 1). Comparing the effects across the factors, repeated measures 4-way ANOVA revealed significant main effects for epoch, light delivery, and group, but not cued side, and included several significant interactions across factors (Table 1). In particular, we found significant interactions between group and all other factors (light delivery, cued side, and delay epoch). Thus, our main finding with optogenetic PS D1-MSN stimulation during the delay period of the task was that light-evoked increases in lick probability were significantly greater during the late delay than the early delay.

**Table 1.** Effects of D1-MSN activation on lick probability (related to Figure 5F-G).

|  | Contra-cued trials |  | Ipsi-cued trials |  |
| --- | --- | --- | --- | --- |
|  | No light | Light | No Light | Light |
| *Mean (s.e.m.)* |  |  |  |  |
| PS ChR2-mCherry (N = 6) |  |  |  |  |
| Early delay | 0.026 (0.004) | 0.129 (0.029) | 0.043 (0.014) | 0.105 (0.031) |
| Late delay | 0.086 (0.026) | 0.533 (0.056) | 0.083 (0.022) | 0.352 (0.053) |
| PS mCherry (N = 4) |  |  |  |  |
| Early delay | 0.037 (0.032) | 0.019 (0.018) | 0.061 (0.017) | 0.044 (0.032) |
| Late delay | 0.064 (0.047) | 0.052 (0.033) | 0.031 (0.017) | 0.084 (0.081) |
| DMS ChR2-mCherry (N = 5) |  |  |  |  |
| Early delay | 0.030 (0.011) | 0.041 (0.007) | 0.064 (0.013) | 0.079 (0.021) |
| Late delay | 0.089 (0.018) | 0.603 (0.080) | 0.100 (0.032) | 0.513 (0.103) |
| DMS mCherry (N = 4) |  |  |  |  |
| Early delay | 0.083 (0.033) | 0.058 (0.013) | 0.044 (0.005) | 0.043 (0.008) |
| Late delay | 0.038 (0.014) | 0.075 (0.026) | 0.066 (0.017) | 0.084 (0.029) |
| | df | F | p | Partial $\eta^2$ |
| *PS 4-way ANOVA* |  |  |  |  |
| Epoch | 1,7 | 48.32 | 0.0001 | 0.93 |
| Light delivery | 1,7 | 59.93 | 0.0001 | 0.94 |
| Group | 1,7 | 24.59 | 0.0011 | 0.95 |
| Cued side | 1,7 | 0.72 | 0.42 | N/A |
| Group x Light | 1,7 | 58.19 | 0.0001 | 0.94 |
| Group x Epoch | 1,7 | 33.45 | 0.0004 | 0.91 |
| Epoch x Light | 1,7 | 45.13 | 0.0001 | 0.89 |
| Group x Light x Cued side | 1,7 | 6.04 | 0.0394 | 0.62 |
| Group x Light x Epoch | 1,7 | 26.10 | 0.0009 | 0.82 |
| Group x Light x Cued side x Epoch | 1,7 | 6.67 | 0.0325 | 0.45 |
| *DMS 4-way ANOVA* |  |  |  |  |
| Epoch | 1,6 | 20.57 | 0.0027 | 0.97 |
| Light delivery | 1,6 | 33.23 | 0.0007 | 0.96 |
| Group | 1,6 | 18.32 | 0.0037 | 0.96 |
| Cued side | 1,6 | 0.03 | 0.87 | N/A |
| Group x Light | 1,6 | 29.48 | 0.001 | 0.96 |
| Group x Epoch | 1,6 | 18.15 | 0.0037 | 0.97 |
| Epoch x Light | 1,6 | 37.53 | 0.0005 | 0.96 |
| Group x Light x Cued side | 1,6 | 1.30 | 0.29 | N/A |
| Group x Light x Epoch | 1,6 | 26.38 | 0.0013 | 0.94 |
| Group x Light x Cued side x Epoch | 1,6 | 0.66 | 0.44 | N/A |

For light delivered to the PS during the orientation-change event, because performance was already at ceiling, it was not possible to observer increases in lick rate. However, we did find effects on lick reaction time. Delivering light to the PS during the change event significantly reduced the median response latency in ChR2-mCherry mice but not in mCherry control mice (Figure 5H; Table 2). A repeated measures 3-way ANOVA showed a significant main effect of light delivery, but not of group nor cued side (Table 2).

**Table 2.** Effects of D1-MSN activation on change event reaction time (ms) (related to Figure 5H)

|  | Contra-cued trials |  | Ipsi-cued trials |  |
| --- | --- | --- | --- | --- |
|  | No light | Light | No Light | Light |
| *Mean (s.e.m.)* |  |  |  |  |
| PS ChR2-mCherry<br>(N = 6) | 333 (19) | 311 (16) | 342 (18) | 323 (20) |
| PS mCherry<br>(N = 4) | 348 (14) | 341 (13) | 361 (8) | 343 (12) |
| DMS ChR2-mCherry<br>(N = 5) | 334 (21) | 320 (18) | 352 (18) | 328 (15) |
| DMS mCherry<br>(N = 4) | 358 (29) | 358 (31) | 337 (24) | 333 (21) |
| | df | F | p | Partial $\eta^2$ |
| *PS 3-way ANOVA* |  |  |  |  |
| Light delivery | 1,8 | 50.27 | 0.0001 | 0.75 |
| Group | 1,8 | 0.78 | 0.40 | N/A |
| Cued side | 1,8 | 2.98 | 0.12 | N/A |
| *DMS 3-way ANOVA* |  |  |  |  |
| Light delivery | 1,7 | 4.90 | 0.06 | N/A |
| Group | 1,7 | 0.21 | 0.66 | N/A |
| Cued side | 1,7 | 0.27 | 0.62 | N/A |

### Comparison with stimulation of the dorsomedial striatum

In the same mice, we also tested the effects of DMS direct pathway stimulation during the task. Our group previously demonstrated that DMS D1-MSN activation biases lick responses in a manner dependent on spatial cues and reward conditions(Wang et al., 2018; Wang and Krauzlis, 2020). Here, to compare with our PS activation, we tested whether temporal expectation also influenced the behavioral effects of DMS direct pathway activation. We found that light delivery to the DMS during the early delay did not significantly change lick probability for either ChR2-mCherry mice or mCherry control mice (Table 1). Light delivery to the DMS during the late delay did significantly increase lick probability in ChR2-mCherry mice but not mCherry control mice (Table 1). As with PS, a repeated measures 4-way ANOVA revealed significant main effects for epoch, light delivery, and group, but not cued side, and included several significant interactions (Table 1). Our group has previously observed an effect of spatial cueing on DMS-evoked licking (Wang et al., 2018; Wang and Krauzlis, 2020). The lack of effect in the present cohort may be due to a smaller group size, as not all mice demonstrate cueing effects, even in the original studies.

We also examined the effects of DMS direct pathway stimulation on the response to the change event. Unlike our results with PS stimulation, we did not find a significant effect of light delivery on median response latency for ChR2-mCherry mice or mCherry mice (Table 2). A repeated measures 3-way ANOVA revealed a trending but insignificant main effect of light delivery (Table 2). Our group has previously demonstrated that activating DMS D1-MSNs significantly reduces event detection latency (Wang et al., 2018); the negative result here may be due to an underpowered cohort size.

Thus, stimulation of either the PS or DMS biased decisions in our visual change-detection task, and both showed larger effects when the stimulation was applied late in the delay period, when the change event was expected to occur. These results show that stimulation of either PS or DMS does not automatically evoke a lick response but instead depends on the context and temporal structure that the mice have learned to associate with rewards in our visual task.

### Effects of stimulating PS depend on the visual context and reward reinforcement

To further elucidate the conditions in which PS direct pathway activation biases visual decisions we performed two additional behavioral experiments in the same cohort. First, we tested whether the induced lick response depended on the presence of choice-relevant visual stimuli (i.e. the Gabor patch). Using our standard visual change-detection task, we introduced a minority of trials in which the stimulation was paired with either the one-patch or pink noise stimuli, but at a time temporally matched to stimulation during the late delay or change epoch (Figure 6A *left*). This experiment replicated our earlier finding that stimulation during contralaterally cued two-patch visual presentation increased the mean lick probability (no light: 0.067 ± 0.034 s.e.m. vs. light: 0.491 ± 0.044). In contrast, pairing PS D1-MSN activation with the contralateral one-patch stimulus resulted in a lick probability of 0.249 ± 0.054, approximately half as much as the two-patch, and light delivery paired with pink noise produced a lick probability of 0.018 ± 0.012, essentially zero (Figure 6A *right*). A repeated measures 1-way ANOVA showed a significant main effect of visual stimulus (F(3,15) = 40.17, P < 0.0001, d = 4.23) on lick probability. Thus, the effects of PS stimulation were not simply due to timing in the task but depended on visual and behavioral context. Evoked-lick rates were highest for the two-patch presentation, which is the context associated with the possibility of a reward-related visual event.

**Figure 6.**
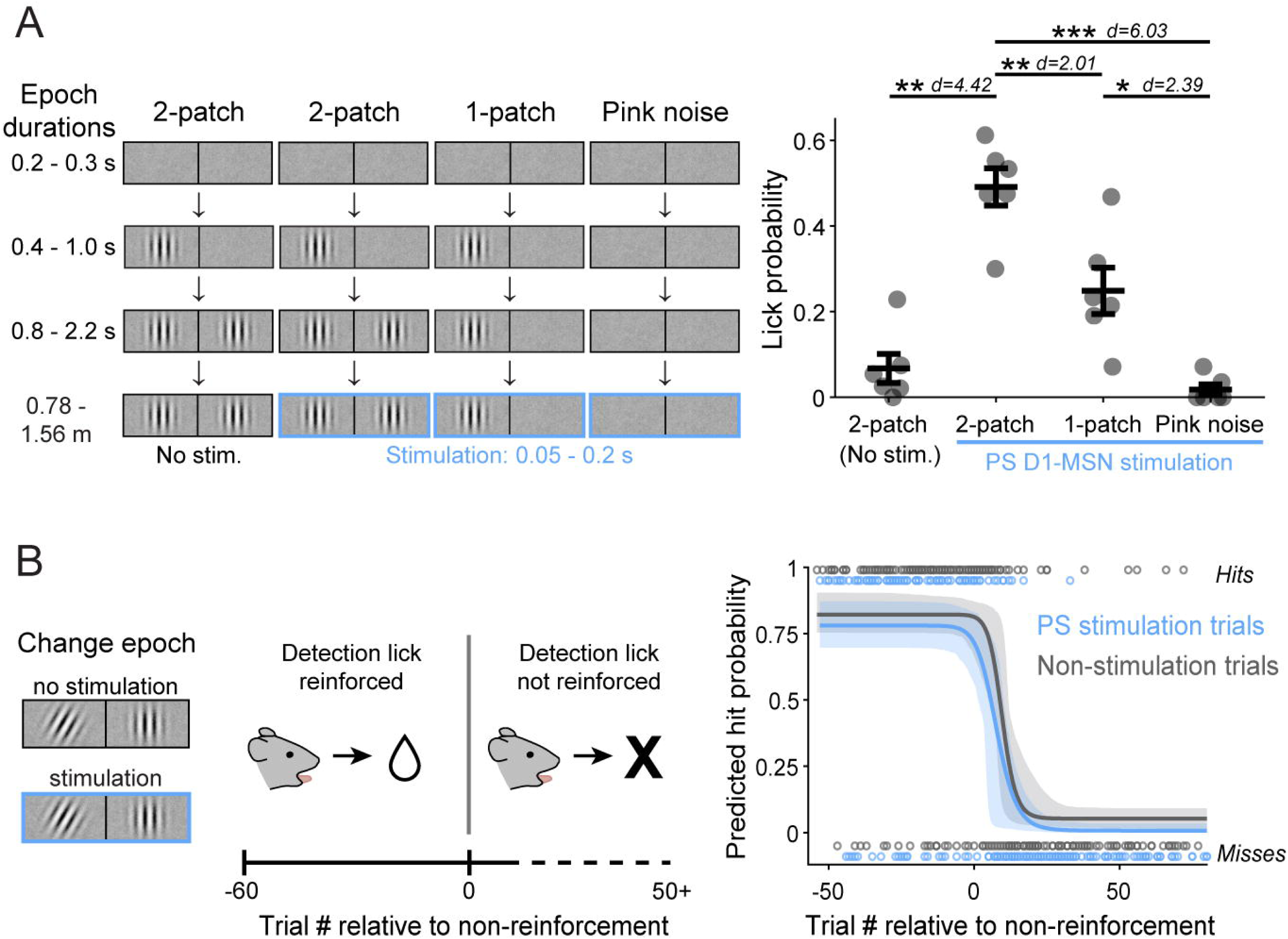
PS direct pathway bias of visual decisions depends on visual context and reward reinforcement. A. Left: Schematic of visual displays and timing of light delivery for two-patch (late delay) and sequence-arrested one-patch and pink noise control trials. Right: Lick probability observed following PS D1-MSN stimulation paired with the different visual conditions on contralaterally cued trials (N = 6 mice). B. Left: Schematic of operant extinction session structure, with change event paired or unpaired with PS D1-MSN stimulation Right: Omitting rewards for correct detection responses resulted in similar extinction rates for PS D1-MSN stimulation (blue) and non-stimulation (gray) trials. Extinction rates derived from trial hits and misses (open circles; combined sessions of 4 mice). One-way repeated measures ANOVA (A). Error bars represent s.e.m. (A); dashed lines represent 95% confidence intervals (CIs) (B).

We next asked whether the decision bias mediated by PS stimulation also depended on reward reinforcement. Specifically, if rewards were withheld for correct responses in our visual task, would PS D1-MSN activation still evoke licks, or otherwise delay the emergence of operant extinction? To test this, we ran mice on a modified version of the task in which only contralaterally cued trials were presented (50% resulted in the change event; 50% of both trial types were paired with PS D1-MSN stimulation). After 60 trials in which correct detection responses were rewarded, trials continued as before but with no rewards administered for the remainder of the session (Figure 6B *left*). Pooling across 10 sessions from 4 mice, we quantified the extinction rates separately for non-stimulation and PS D1-MSN stimulation trials by fitting the hits and misses relative to the first non-rewarded trial in a four-parameter logistic regression model with bootstrapped computed 95% confidence intervals (CIs) (Figure 6B *right*). We found that the fitted model curve for stimulation trials did not significantly differ from that of non-stimulation trials. A predicted hit rate of 0.5 occurred at 8.9 trials (95% CIs: 5, 11.8) post-reward non-reinforced on the curve fitted to the non-stimulation trials and 5.8 trials (CIs: 1.4, 9.1) on the curve fitted to stimulation trials (z-test z = -1.673, P = 0.094). Thus, PS stimulation did not change the time course of operant extinction, and the effects of PS stimulation disappeared as extinction took hold.

### The basal ganglia direct pathway modulates superior colliculus neuronal activity during visual decision-making

Having demonstrated that both PS and DMS D1-MSN activation robustly affects visual decisions in our task, we next investigated one of the major output targets of these basal ganglia circuits and how neuronal activity was modulated during the visual detection task. The SC is a shared target of both PS- and DMS-originating direct pathways (Figure 2K-R, Figure 3C-F) and plays a crucial role in visual processing and visually guided decisions in rodents (Cang et al., 2018; Wang et al., 2020). We collected extracellular electrophysiological recordings in the SC of our cohort of PS and DMS D1-MSN ChR2-mCherry expressing mice (N = 6) performing the orientation change-detection task with and without optogenetic stimulation. PS or DMS D1-MSN activation was tested in separate recording sessions.

We recorded a total of 530 single units across the layers of the SC (Figure 7A-B). Grouping these units by their estimated location in either superficial (sSC) or deeper (dSC) layers, we observed neuronal responses to key features of the spatially cued change-detection task (Figure 7C-D; Table 3). Phasic responses to the onset of the cue patch, when presented in the contralateral visual field, were present in the majority of SC neurons: 78.4% of sSC units and 55.7% of dSC units. Delay activity (defined as a significant difference in firing rates in the late vs early delay epochs) was also observed in many neurons: 46.6% of sSC units and 37.3% of dSC units. We observed spatial cue-related modulation (defined as the difference in firing rate between contra- and ipsi-laterally cued trials late in the delay epoch) but this was present in a minority of neurons: 18.2% of sSC units and 20.6% of dSC units. Similarly, phasic responses to the orientation change were observed in 18.2% of sSC units and 22.4% of dSC units. Lastly, we found lick-related activity (defined as the difference in firing rate in a 100 ms window centered on the first detection lick compared to pre-change event firing activity) in a small minority of SC neurons: 1.1% of sSC units and 6.6% of dSC units. Comparing the modulation between the superficial and deep layers, we found that sSC units were significantly more likely to have a cue response than dSC units (χ^2^ = 15.76, P < 0.0001, d = 0.35) whereas dSC units were significantly more likely to have lick-related activity (χ^2^ = 4.04, P = 0.044, d = 0.18).

**Figure 7.**
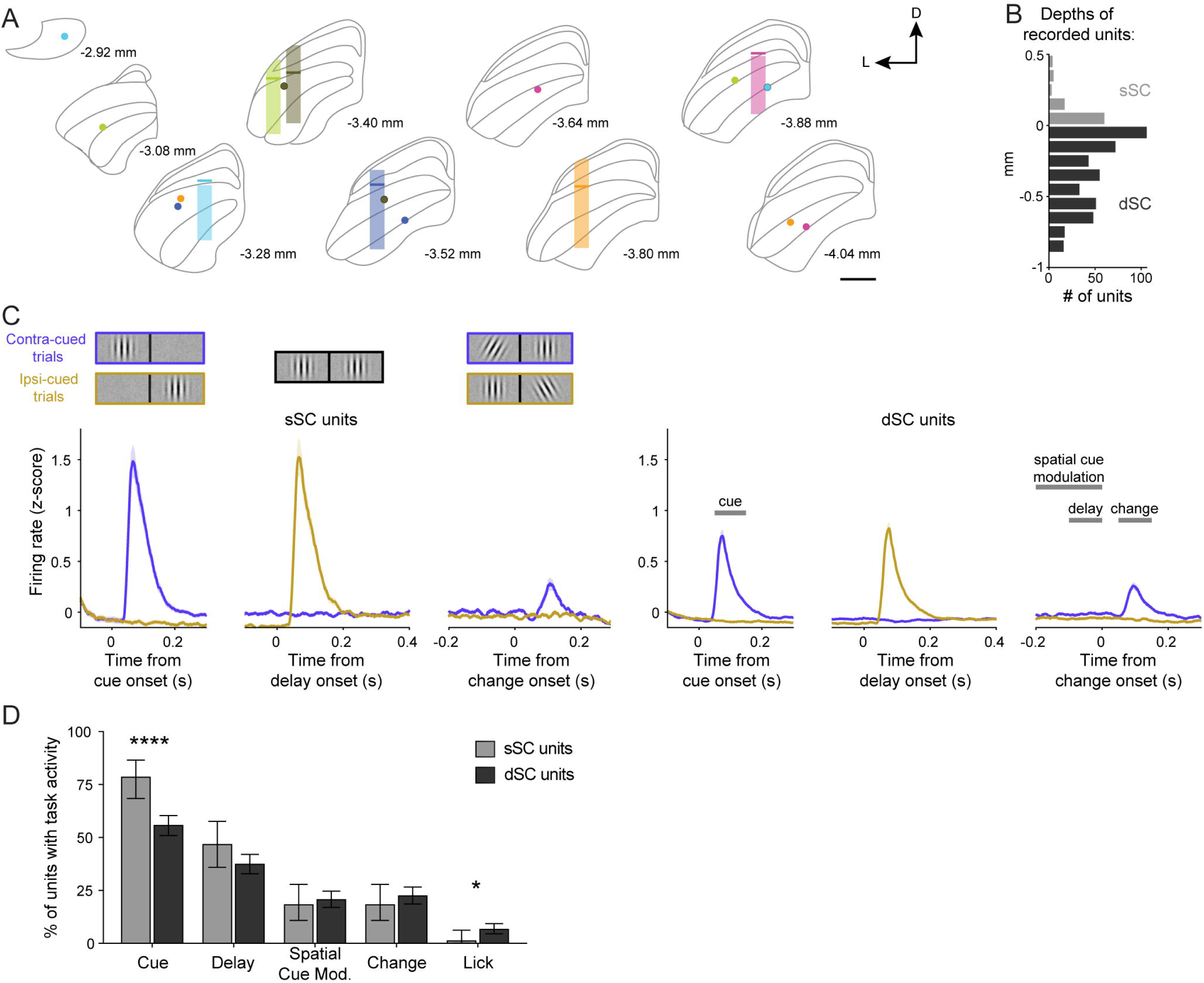
PS and DMS direct pathways preferentially modulate SC neurons responsive to visual task properties. A. Approximated regions of the SC recorded from 6 mice on a standard atlas (Franklin and Paxinos, 2008). Transparent bar represents the anterior-posterior and medial-lateral location of the recording bundle center, with approximate DV range of recorded units relative to the division between superficial (sSC) and deeper (dSC) layers (solid bar). Filled circles indicate location of electrolytic lesioned wire tips, color-coded per mouse. B. Distribution of recorded single units across the SC relative to the estimated division of sSC and dSC layers (n = 530 units; N = 6 mice). C. Left: PSTHs of sSC single unit responses to the spatially cued orientation change task stimuli (n = 88). Right: PSTHs of dSC single unit responses with time windows for quantifying task-related activity (n = 442). D. Percentages of sSC (light gray) and dSC (dark gray) units that exhibited task-related activity. Chi-square tests (D). Error bars represent s.e.m. (C) or 95% CI (D). Scale bar = 500 um. Abbreviations: D (dorsal), L (lateral).

**Table 3.** Percentages (95% CIs) of SC units modulated by task properties and D1-MSN activation (related to Figure 7D and Figure 8C).

|  | Cue<br>Event | Delay<br>Activity | Spatial Cue<br>Mod. | Change<br>Event | Lick<br>Activity |
| --- | --- | --- | --- | --- | --- |
| All sSC units | 78.4<br>(68.4, 86.5) | 46.6<br>(35.9, 57.5) | 18.2<br>(10.8, 27.8) | 18.2<br>(10.8, 27.8) | 1.1<br>(0.03, 6.2) |
| All dSC units | 55.7<br>(50.9, 60.4) | 37.3<br>(32.8, 42.0) | 20.6<br>(16.9, 24.7) | 22.4<br>(18.6, 26.6) | 6.6<br>(4.4, 9.3) |
| PS mod. &<br>Task non-mod.<br>dSC units | 21.3<br>(12.7, 32.3) | 21.9<br>(14.7, 30.7) | 22.9<br>(16.2, 30.7) | 19.9<br>(13.6, 27.4) | 27.6<br>(21.2, 34.6) |
| PS mod. &<br>Task mod. dSC<br>units | 31.5<br>(23.0, 41.0) | 36.1<br>(25.1, 48.3) | 41.3<br>(27.0, 56.8) | 51.1<br>(35.8, 66.3) | 0<br>(0, 97.5) |
| DMS mod. &<br>Task non-mod.<br>dSC units | 36.1<br>(26.6, 46.5) | 29.0<br>(21.7, 37.1) | 31.4<br>(24.9, 38.5) | 29.4<br>(22.9, 36.7) | 38.9<br>(32.3, 45.9) |
| DMS mod. &<br>Task mod. dSC<br>units | 36.6<br>(28.4, 45.5) | 49.4<br>(38.2, 60.6) | 62.2<br>(47.4, 76.1) | 62.5<br>(47.4, 76.1) | 10.0<br>(1.2, 31.7) |

We next examined how optogenetic direct pathway activation modulated ipsilateral SC neuronal activity. We classified units as modulated by D1-MSN activation if there was a significant difference in firing during the 150-ms light-delivery period compared to an equivalent window during non-stimulation trials. Because the basal ganglia direct pathway reaches the SC by a series of two inhibitory projections, we expected activation of D1-MSNs to have a predominately excitatory effect on SC neuronal activity (Hikosaka and Wurtz, 1983; Hikosaka et al., 2000; Kaneda et al., 2008; Freeze et al., 2013). Indeed, the majority of units modulated by PS or DMS direct pathway activation showed excitatory changes in firing rate (Figure 8A-B *left*). However, we also observed populations of units that were exclusively inhibited by direct pathway activation (Figure 8A-B *middle*). This effect was more prevalent for PS. Stimulation induced an inhibitory effect in a larger percentage of dSC units modulated by PS (27.5%; 15.9, 41.7 95% CI) than modulated by DMS (13.3%; 6.8, 22.5; χ^2^ = 4.20, P = 0.04, d = 0.36). Finally, we observed a small number of units that demonstrated both excitation and inhibition, either in different test epochs or within the same epoch (Figure 8A-B *right*).

**Figure 8.**
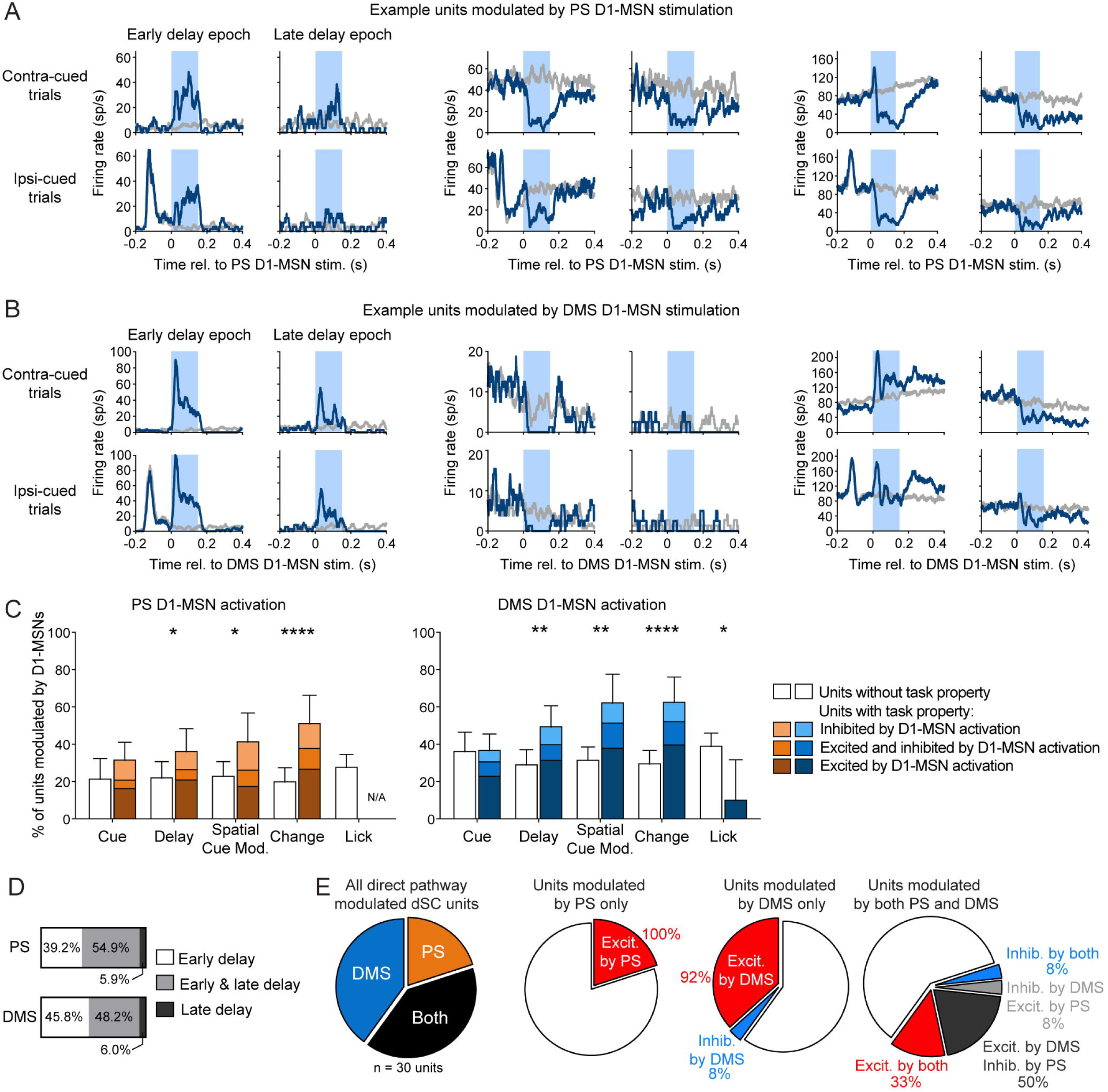
Superior colliculus neuronal responses to PS and DMS direct pathway activation. A. Mean firing rates for example dSC units that were excited (left), inhibited (center), or both excited and inhibited (right) by PS D1-MSN optogenetic stimulation (blue) compared to activity on non-stimulation trials (gray). Optogenetic light delivery indicated by light blue shading. B. Same as A but for dSC units modulated by DMS D1-MSN stimulation. C. Percentages of dSC units exhibiting task properties (colored bars) or not (white bars) that were modulated by optogenetic activation of PS (left) or DMS (right) D1-MSNs. D. Proportions of PS (top) and DMS (bottom) D1-MSN modulated dSC units that were modulated during early delay (white), late delay (dark gray), or both epochs (light gray). E. Left: from recording sessions with both DMS and PS D1-MSN stimulation, proportions of dSC single units that were modulated by PS, DMS or both D1-MSN pathways. Center and right: the type of modulatory effect on dSC firing by PS and DMS D1-MSN stimulation.

We examined how SC units that exhibited different task-related properties were modulated by direct pathway stimulation. For the sSC, we found very few units to be modulated by either PS (4 out of 35 units) or DMS (3 out of 45 units) D1-MSN activation. Therefore, we restricted our analyses to dSC units, among which 51 out of 186 units were modulated by PS direct pathway activation and 83 out of 228 units were modulated by DMS direct pathway activation. We observed that activation of the striatal direct pathways did not affect all dSC units equally. For the PS, we found that D1-MSN activation was more likely to modulate units that exhibited delay activity (χ^2^ = 4.46, P = 0.035, d = 0.31), showed spatial cue-related modulation (χ^2^ = 5.92, P = 0.015, d = 0.36), or responded to the orientation change (χ^2^ = 16.74, P < 0.0001, d = 0.63) (Figure 8C *left*; Table 3). Similarly, for the DMS, D1-MSN activation preferentially modulated units with delay activity (χ^2^ = 9.52, P = 0.002, d = 0.42), spatial cue-related modulation (χ^2^ = 12.66, P = 0.0004, d = 0.48), or that responded to the orientation change (χ^2^ = 17.89, P < 0.0001, d = 0.58). DMS D1-MSN activation was also more likely to modulate units that did not show lick related activity (χ^2^ = 6.60, P = 0.0102, d = 0.35) (Figure 8C *right*; Table 3). However, we note that we had relatively few units with lick-related activity in our recordings, likely because most of our recording sites were in the central SC, rather than the lateral SC, where lick-related units tend to be concentrated (Thomas et al., 2023).

We also examined whether dSC modulation by direct pathway activation depended on the stimulated epoch. For PS stimulation, we found that approximately half of D1-MSN modulated dSC units were modulated during both the early and late delay epochs (54.9%), whereas 39.2% were exclusively modulated during the early delay and only a small percentage were modulated exclusively during the late delay (5.9%) (Figure 8D). dSC units modulated by DMS D1-MSN stimulation showed similar distributions: 48.2% for both epochs, 45.8% exclusively during the early delay, and 6.0% exclusively during the late delay (Figure 8D).

### Direct comparison of PS and DMS activation on SC unit activity

We also were able to directly compare the effects of PS and DMS direct pathway activation on SC units. This comparison at the neuronal level is of particular interest because our histological tracing of PS and DMS direct pathways targeting the SC showed distinct but overlapping patterns of axonal innervation in the SC. We investigated whether these pathways target the same neuronal populations within the SC. In two mice, we recorded from SC units while stimulating PS or DMS D1-MSNs during the early delay in the same session. We recorded a total of 30 optogenetically modulated single units: 40% were exclusively modulated by DMS direct pathway activation, 20% were exclusively modulated by PS direct pathway activation, and 40% were modulated by both pathways (Figure 8E). Of the units that were modulated by both direct pathways, 58.3% exhibited different modulatory effects, that is, these units were excited by one pathway and inhibited by the other. This finding suggests that the PS and DMS direct pathways may innervate different SC populations.

## Discussion

The present study investigated the contributions of the mouse PS to visual behavior. Our results demonstrate that the PS is uniquely anatomically positioned to relay visual signals through the basal ganglia. Compared to the DMS, which has been studied more frequently in visual tasks, the PS is more strongly innervated by visual cortex and gives rise to an alternative direct pathway for visual processing. Our results show that activating this pathway robustly and selectively modulates visual decisions. We also identified that both PS and DMS direct pathway activation targeted neurons in the SC that were sensitive to context and reward-associated stimuli. Together, these findings further demonstrate distinct circuits through the basal ganglia for visual perceptual decision-making.

The effects of PS direct pathway manipulation on visual decisions were similar to that of the DMS (Wang et al., 2018), suggesting a comparable influence on perceptual behavior, at least as revealed in our visual task. Similar to the DMS, we found that stimulating the PS direct pathway does not drive an obligatory and fixed lick response. Optogenetic activation induced ‘yes’ perceptual responses that depended on visual and temporal context. Moreover, PS D1-MSN stimulation did not delay extinction of the response to the devalued change event when rewards were withheld. These findings are consistent with the view that the direct pathway provides a permissive gating function for actions driven by circuits extrinsic to the basal ganglia (Hormigo et al., 2021). In the context of perceptual decision-making, the convergence of cortical and subcortical inputs to the striatum is proposed to similarly enable the selection and implementation of visual-based decision policies (Ding, 2023).

The PS direct pathway has also been shown to bias auditory-based decisions (Guo et al., 2018; Chen et al., 2022). Dopaminergic signaling in this region reinforces the avoidance of novel or threatening stimuli of multiple sensory modalities (Menegas et al., 2018) and is also shown to scale with auditory uncertainty (Chen et al., 2022). Moreover, activating PS D1-MSNs in freely moving mice failed to induce orienting responses which are reliably observed with dorsal striatal direct pathway activation (Guo et al., 2018). The combination of sensitivity to multimodal stimuli and distinct dopaminergic activity positions the PS to uniquely contribute to decision-making and behavior.

Although PS and DMS stimulation yielded comparable behavioral effects in our paradigm, differences in the connectivity of these striatal subregions hint at potential functional specializations, starting with the sources of their cortical afferents. The DMS was preferentially innervated by anterior primary and secondary visual areas. Neurons in mouse anterior secondary areas are noted to preferentially respond to stimuli of lower spatial and higher temporal frequencies, exhibit stronger orientation selectivity, and prefer fast-moving stimuli (Han et al., 2022). In contrast, we found that the PS was innervated by all of secondary visual cortex, including posterior areas that are characterized by a preference for higher spatial, lower temporal, and slow-moving stimuli (Han et al., 2022). Whether this broader anatomical connection confers the PS a relative sensitivity or contextual selectively for visual input remains to be determined. Differences in striatal afferents extend beyond the visual cortex. Whereas the DMS is preferentially innervated by frontal cortices including anterior cingulate, prelimbic and orbital cortices, the PS notably receives input from all sensory cortical areas (Hintiryan et al., 2016; Hunnicutt et al., 2016; Lee et al., 2023).

There are also differences in the efferent pathways from the PS and DMS, giving rise to distinct circuits through the basal ganglia that target different topographical subregions of the output nuclei (Lee et al., 2020; Miyamoto and Fukuda, 2022). The DMS direct pathway is routed through the SNr and EP, whereas the PS direct pathway innervates the SNl and PP. Although the specific functions of the SNl and PP are not yet well defined, our findings of robust behavioral effects induced by PS D1-MSN activation indicate that these nuclei should be considered among the basal ganglia nuclei involved in perceptual decision-making. Previous studies of the SNl and PP that have examined the anatomical properties of these nuclei suggest a range of possible circuit mechanisms. The SNl consists of dopaminergic and GABAergic neurons that differentially project to sensory nuclei, motor-related regions, the amygdala, as well as forming loops back to the basal ganglia (Kaelber and Afifi, 1979; Moriizumi et al., 1992; Takada, 1992; Todd et al., 2022). The adjacent PP innervates a range of midbrain, cortical, and diencephalon nuclei (Arnault and Roger, 1987; Cai et al., 2024).

Given that the PS and DMS route through different output nuclei, it is not surprising that these two striatal regions yield divergent efferent connectivity in the thalamus and midbrain (Benavidez et al., 2021; McElvain et al., 2021). We found marked differences in the innervation of one prominent target, the SC. Compared to the DMS-originating direct pathways, the PS to SNl/PP circuit exhibited preferential innervation of the lateral SC, which is notably connected with cortical, thalamic, and hindbrain nuclei associated with visuomotor and somatic sensorimotor functions (Benavidez et al., 2021). In contrast, the direct pathways initiated from the DMS did not show medial-lateral differences in SC innervation, and this more uniform innervation of SC deeper layers suggests engagement with additional anatomical networks including those involving the dorsal lateral geniculate nucleus as well as secondary visual and association cortical areas (Benavidez et al., 2021).

Related to these different efferent circuit projections, we also identified differences in how activation of the direct pathways alters neuronal activity in the SC. We observed that PS direct pathway activation induced inhibition in a relatively larger population of dSC units. Given the established disinhibitory effect of the direct pathway, excitation was the expected, and indeed, predominate modulatory effect on SC unit firing. However, suppression may occur if the recorded SC unit is efferent to a local interneuron that is itself disinhibited by the direct pathway resulting in a tri-synaptic inhibitory configuration (e.g., D1-MSN→SN/EP→SC interneuron→recorded SC unit). A larger proportion of SC units inhibited by PS D1-MSN activation suggests that the innervation of the SC by this direct pathway may be more biased toward interneurons.

We found that both PS and DMS direct pathways preferentially modulated dSC units with specific types of task-related activity. Units that exhibited delay-related activity, spatial cue-related modulation, or phasically responded to the reward-associated orientation change were more likely to be modulated by direct pathway activation. Interestingly, we did not find a similar preference for units that responded to the appearance of the Gabor patch cue. This suggests that the direct pathway bias is not merely for visually responsive SC units, but for those processing aspects of visual signals related to a goal or outcome. Our group has previously shown that dSC cue-related modulation contributes to spatial attention by biasing signal processing for reward-associated event locations (Wang et al., 2022). Preferential innervation of neurons showing attention-like activity may be one mechanism by which the direct pathway biases decisions in manner linked to the allocation of spatial attention (Wang and Krauzlis, 2020).

In summary, the results of the present study provide the first evidence for involvement of the mouse PS in visual behavior. Here, we monitored the neuronal modulation of one target of the basal ganglia direct pathway, the SC, during a visual decision-making task. In addition to the several circuit routes comprising the direct pathway that we have already noted, there are also many output targets that may contribute to perceptual decisions. The basal ganglia direct pathways also target the motor thalamus, reticular formation, and pedunculopontine nucleus, which may each also contribute to the behavioral response (van der Kooy and Carter, 1981; McElvain et al., 2021). Although it remains an open challenge to understand how these complex circuits operate together for visual perception, our results show that any description should include circuits that originate in the posterior striatum.

## Acknowledgements

This work is supported by the National Eye Institute Intramural Research Program at the National Institutes of Health ZIA EY000511 (R.J.K.). The authors thank Dr. Carlos Mejias-Aponte for assistance with histological analysis.

